# Persistent circulation of Rift Valley fever virus lineage C in Rwanda, 2022–2025

**DOI:** 10.64898/2026.06.04.730228

**Authors:** Jean Claude Udahemuka, Hayley Cassidy, Leonard Schuele, Evodie Uwibambe, Methode Gasana Ngabo, Leandre Murhula Masirika, Saria Otani, Misbah Gashegu, Jean-Claude Twizere, Frank Aarestrup, Fabrice Ndayisenga, Bas B. Oude Munnink, Marion P.G. Koopmans, Pacifique Ndishimye

**Affiliations:** Department of Veterinary Medicine, University of Rwanda, Nyagatare, Rwanda; Stansile Research Organization, Kigali, Rwanda & Halifax, Canada; Department of Viroscience, Pandemic and Disaster Preparedness Centre, Erasmus University Medical Center, Rotterdam, the Netherlands; Department of Animal Resource Research and Technology Transfer, Rwanda Agriculture and Animal Resources Development Board (RAB), Huye P.O. Box 5016, Rwanda; Congo Outbreaks, Research for Development (CORD), South-Kivu, Bukavu, Democratic Republic of the Congo; Centre de Recherche en Sciences Naturelles de Lwiro, South-Kivu, Bukavu, Democratic Republic of the Congo; Research Group for Genomic Epidemiology, National Food Institute, Technical University of Denmark, Lyngby, Denmark; Rwanda Biomedical Centre, Kigali, Rwanda; Université de Liège, Liege, Belgium; Epidemic Response Laboratory, African Institute for Mathematical Sciences (AIMS), Kigali, Rwanda

**Keywords:** Rift Valley fever virus, Whole-genome sequencing, Metagenomics, Nanopore sequencing, Public Health Surveillance, Genomics

## Abstract

Rwanda has experienced recurrent Rift Valley fever virus outbreaks in the last decade. In this study, we investigated whether these outbreaks resulted from repeated introductions or sustained local circulation. We generated RVFV whole-genome sequences from livestock samples collected between 2022 and 2025 using Nanopore sequencing. Genomic analyses indicated the outbreaks resulted from sustained local circulation of lineage C rather than repeated introductions, suggesting ongoing transmission likely driven by sporadic spillover. This study underscores the importance of continuous genomic surveillance in endemic settings.

## Introduction

Rift Valley fever virus (RVFV) is a significant threat to human and animal health across East and Sub-Saharan Africa with increasing frequency and scale of outbreaks (1). This is underscored by the recent 2025 RVFV outbreak in Senegal and Mauritania (2), affecting both humans and animals, with ongoing transmission reported as of January 2026 (3,4). Transmission typically occurs through Aedes, and to a lesser degree Culex mosquito bites or contact with contaminated blood, primarily affecting cattle, sheep, and goats (5). This can result in fever, hemorrhaging and “abortion storms” leading to significant economic losses (6). Humans can get infected through mosquito exposure, as well as handling of animal tissues and consumption of unpasteurized milk.

RVFV was first detected in Rwanda in 2012 and has since caused small annual outbreaks, culminating in larger outbreaks in 2022 and 2024 (7,8), with further case numbers in 2025, affecting different regions within Rwanda. It is unknown whether the different outbreaks are linked to a single introduction or reflect repeated introductions or both. Here, we present the genomic characterization of RVFV from livestock with initial genome detection through metagenomic analysis from samples collected during the 2024 outbreak, and subsequent expansion of the study to include a wider representation of samples collected between 2022 and 2025 using real-time field-based Nanopore sequencing.

## Materials and Methods

Blood samples were collected from suspected livestock cases (cattle, sheep and goats) by the Rwanda Agriculture and Animal Resources Development Board (RAB) and screened for RVFV using RealStar® RVFV RT-PCR Kit 1.0 (Altona Diagnostics). Positive blood samples were then selected for Nanopore sequencing during two capacity building workshops in Tanzania and Rwanda in October 2024 and 2025 respectively, with local follow-up sequencing onsite in Rwanda.

In October 2024, ten RVFV PCR-positive samples collected from livestock during the 2024 outbreak were randomly selected and made available by the Rubirizi Veterinary Virology Laboratory within RAB for metagenomic sequencing as described previously (10). Following the first metagenomic sequencing results, a pan-amplicon scheme was designed using PrimalScheme v3.2.3 (https://github.com/aresti/primalscheme), along with reference genomes (>90% segment coverage) obtained from the BV-BRC database (Table S1, Supplementary Materials). The design produced 65 overlapping primers with 400bp amplicon length in two separate pools (Table S2, Figure S1). In October 2025, retrospective samples collected from livestock during the 2022 (n= 14) and 2024 outbreaks (n=1), along with four samples collected in 2025 were selected for amplicon sequencing (Supplementary Materials). This selection was done based on sample availability. Sequencing was performed on site using the MinION device and basecalling was performed using Dorado v7.9.8.

## Results and Discussion

RVFV surveillance in Rwanda between 2022-2025 indicated low overall prevalence, potentially reflecting limited testing, but increasing positivity over time, with outbreaks affecting multiple livestock species across diverse regions (Table S3, Figure 1).

**Figure 1.**
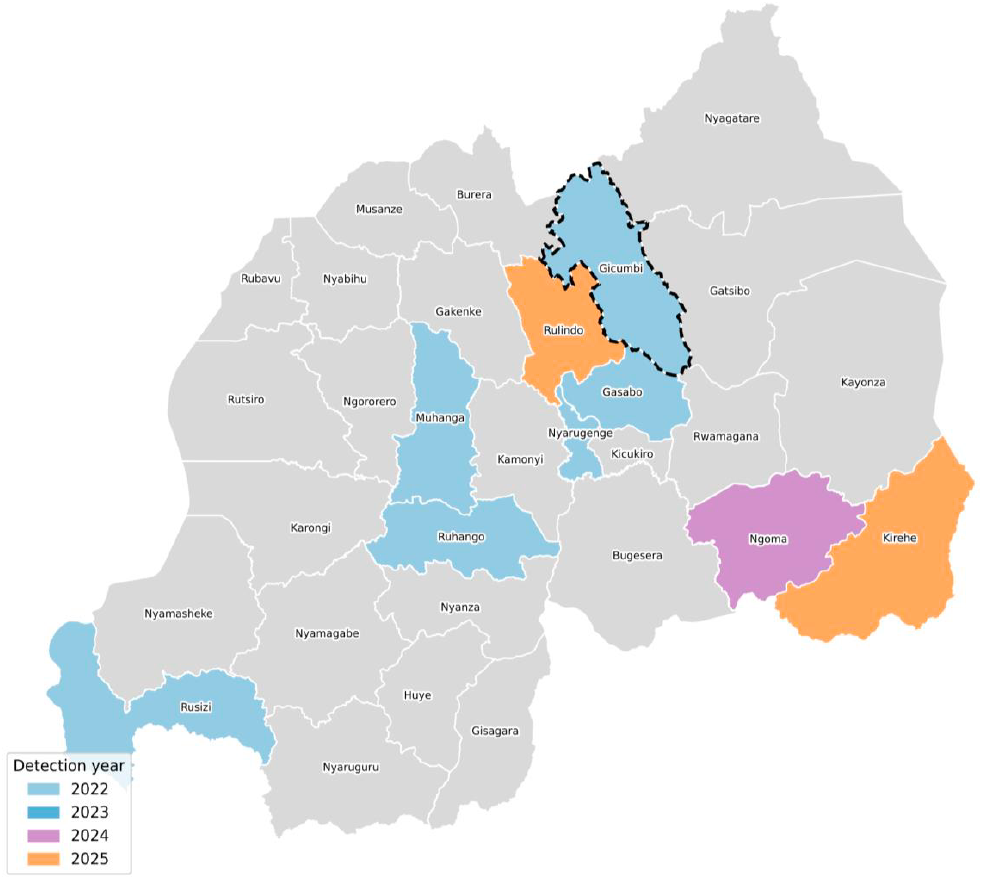
RVFV detection by district in Rwanda between 2022-2025. Following positive laboratory confirmation in 2024 in cattle, sheep and goats, Rwanda Agriculture and Animal Resources Development Board reported a RVF outbreak and established several immediate measures to prevent the spread of the disease to other regions (8). These measures included mass testing and vaccination efforts. A total of 111,170 animals were vaccinated in 2024 (35,089 cattle, 73,795 goats, and 2,286 sheep) increasing to 1,484,133 animals (1,051,857 cattle, 368,847 goats and 63,429 sheep) in 2025 using the thermostable Clone 13 vaccine (the total number of vaccinations per district, the total number of farms per district and total number of animals vaccinated per farm was not available) - Personal communication from the Chief Veterinary Officer, National Veterinary Reference Laboratory in Rubirizi, Rwanda, December 2025. The dashed line indicates multiple detection years with RVFV identified in Gicumbi in 2022 and 2023. A more detailed overview of RVFV cases in Rwanda between 2022-2025, including districts tested and the percent positivity is represented in Table S1.

In total, we generated nine near-complete sequences (>95%) from bovine samples using either metagenomic or amplicon-based sequencing, five of which were from 2022, one from 2024 and three from 2025 (Table 1). Among the partial genomes, we generated 18 near-complete L segments, 11 near-complete M segments and 8 near-complete S segments. Overall, amplicon sequencing generated a coverage percentage of 53.65-98.08 (Segment S), 83.6-100% (Segment M) and 63.68-99.91% (Segment L) from samples with varying viral loads (Ct 15.8-30.8)(Figure S2). We did observe one amplicon dropout in Segment M (299bps at approx. position 2188-2487) and one in Segment S (570bps at approx. position 549-1119)(Figure S3).

**Table 1:**
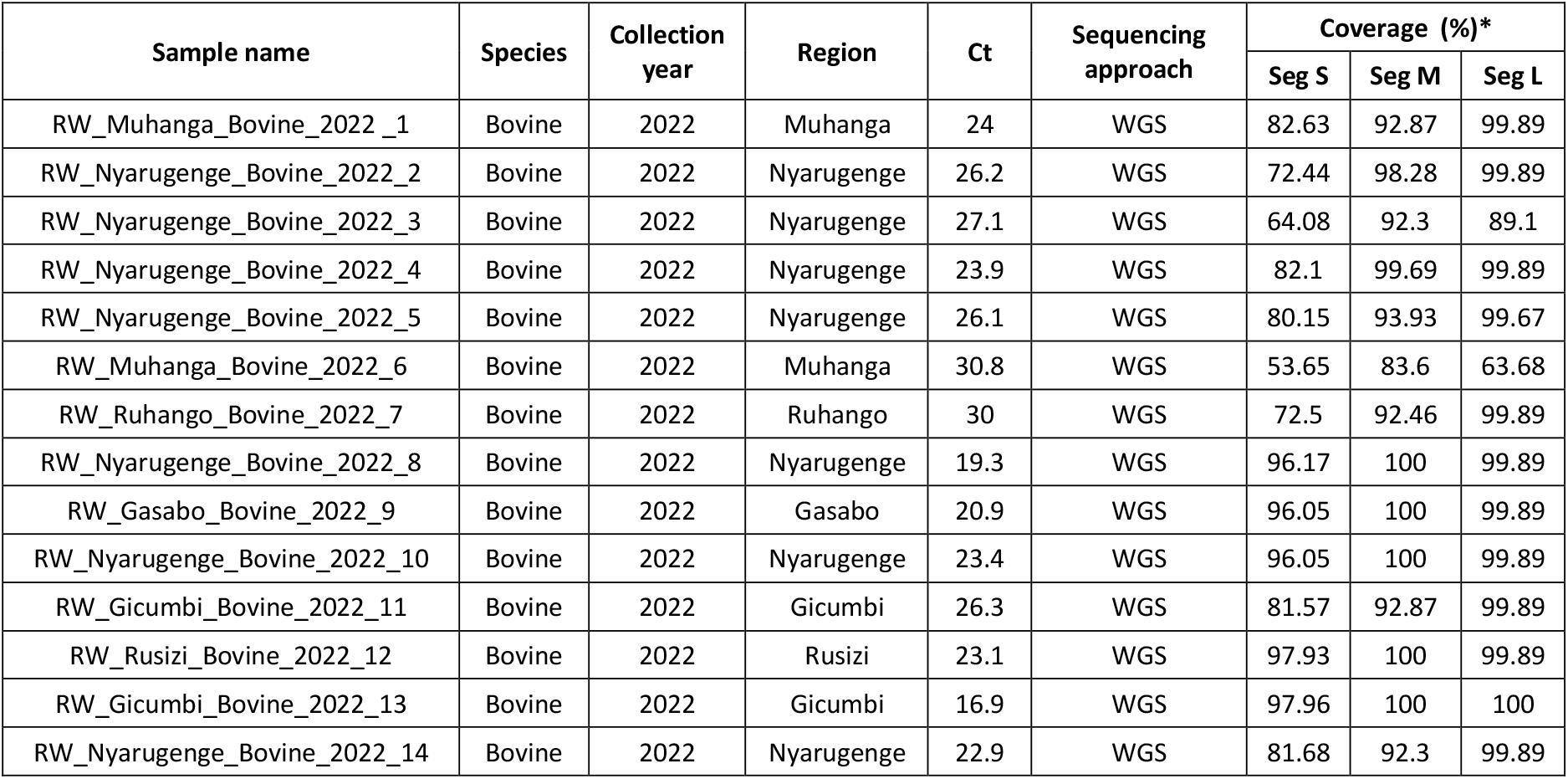
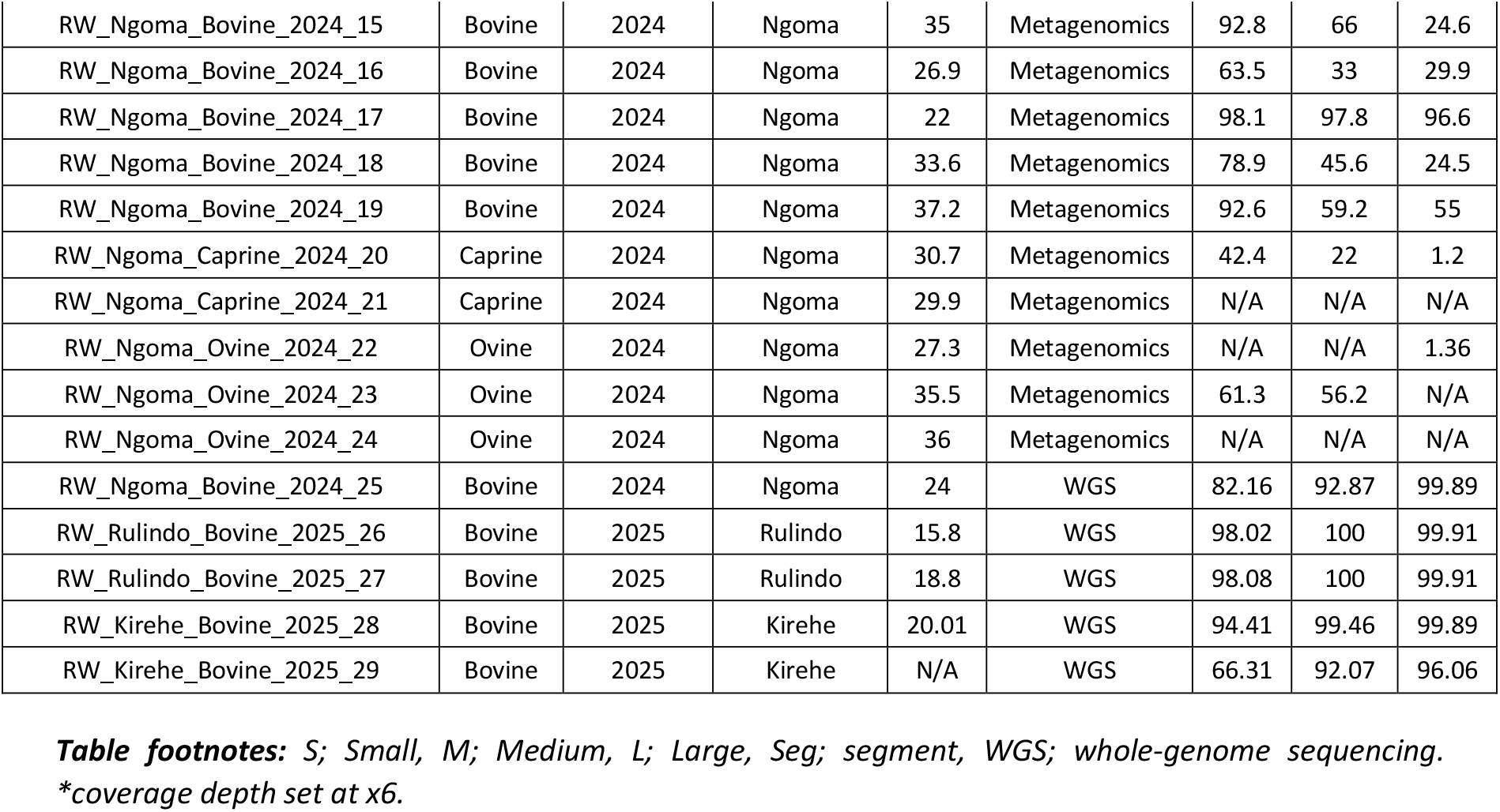
Overview of selected RVFV cases (2022-2025) using two sequencing approaches.

| Sample name | Species | Collection year | Region | Ct | Sequencing approach | Coverage (%)* |  |  |
| --- | --- | --- | --- | --- | --- | --- | --- | --- |
|  |  |  |  |  |  | Seg S | Seg M | Seg L |
| RW_Muhanga_Bovine_2022_1 | Bovine | 2022 | Muhanga | 24 | WGS | 82.63 | 92.87 | 99.89 |
| RW_Nyarugenge_Bovine_2022_2 | Bovine | 2022 | Nyarugenge | 26.2 | WGS | 72.44 | 98.28 | 99.89 |
| RW_Nyarugenge_Bovine_2022_3 | Bovine | 2022 | Nyarugenge | 27.1 | WGS | 64.08 | 92.3 | 89.1 |
| RW_Nyarugenge_Bovine_2022_4 | Bovine | 2022 | Nyarugenge | 23.9 | WGS | 82.1 | 99.69 | 99.89 |
| RW_Nyarugenge_Bovine_2022_5 | Bovine | 2022 | Nyarugenge | 26.1 | WGS | 80.15 | 93.93 | 99.67 |
| RW_Muhanga_Bovine_2022_6 | Bovine | 2022 | Muhanga | 30.8 | WGS | 53.65 | 83.6 | 63.68 |
| RW_Ruhango_Bovine_2022_7 | Bovine | 2022 | Ruhango | 30 | WGS | 72.5 | 92.46 | 99.89 |
| RW_Nyarugenge_Bovine_2022_8 | Bovine | 2022 | Nyarugenge | 19.3 | WGS | 96.17 | 100 | 99.89 |
| RW_Gasabo_Bovine_2022_9 | Bovine | 2022 | Gasabo | 20.9 | WGS | 96.05 | 100 | 99.89 |
| RW_Nyarugenge_Bovine_2022_10 | Bovine | 2022 | Nyarugenge | 23.4 | WGS | 96.05 | 100 | 99.89 |
| RW_Gicumbi_Bovine_2022_11 | Bovine | 2022 | Gicumbi | 26.3 | WGS | 81.57 | 92.87 | 99.89 |
| RW_Rusizi_Bovine_2022_12 | Bovine | 2022 | Rusizi | 23.1 | WGS | 97.93 | 100 | 99.89 |
| RW_Gicumbi_Bovine_2022_13 | Bovine | 2022 | Gicumbi | 16.9 | WGS | 97.96 | 100 | 100 |
| RW_Nyarugenge_Bovine_2022_14 | Bovine | 2022 | Nyarugenge | 22.9 | WGS | 81.68 | 92.3 | 99.89 |
| RW_Ngoma_Bovine_2024_15 | Bovine | 2024 | Ngoma | 35 | Metagenomics | 92.8 | 66 | 24.6 |
| RW_Ngoma_Bovine_2024_16 | Bovine | 2024 | Ngoma | 26.9 | Metagenomics | 63.5 | 33 | 29.9 |
| RW_Ngoma_Bovine_2024_17 | Bovine | 2024 | Ngoma | 22 | Metagenomics | 98.1 | 97.8 | 96.6 |
| RW_Ngoma_Bovine_2024_18 | Bovine | 2024 | Ngoma | 33.6 | Metagenomics | 78.9 | 45.6 | 24.5 |
| RW_Ngoma_Bovine_2024_19 | Bovine | 2024 | Ngoma | 37.2 | Metagenomics | 92.6 | 59.2 | 55 |
| RW_Ngoma_Caprine_2024_20 | Caprine | 2024 | Ngoma | 30.7 | Metagenomics | 42.4 | 22 | 1.2 |
| RW_Ngoma_Caprine_2024_21 | Caprine | 2024 | Ngoma | 29.9 | Metagenomics | N/A | N/A | N/A |
| RW_Ngoma_Ovine_2024_22 | Ovine | 2024 | Ngoma | 27.3 | Metagenomics | N/A | N/A | 1.36 |
| RW_Ngoma_Ovine_2024_23 | Ovine | 2024 | Ngoma | 35.5 | Metagenomics | 61.3 | 56.2 | N/A |
| RW_Ngoma_Ovine_2024_24 | Ovine | 2024 | Ngoma | 36 | Metagenomics | N/A | N/A | N/A |
| RW_Ngoma_Bovine_2024_25 | Bovine | 2024 | Ngoma | 24 | WGS | 82.16 | 92.87 | 99.89 |
| RW_Rulindo_Bovine_2025_26 | Bovine | 2025 | Rulindo | 15.8 | WGS | 98.02 | 100 | 99.91 |
| RW_Rulindo_Bovine_2025_27 | Bovine | 2025 | Rulindo | 18.8 | WGS | 98.08 | 100 | 99.91 |
| RW_Kirehe_Bovine_2025_28 | Bovine | 2025 | Kirehe | 20.01 | WGS | 94.41 | 99.46 | 99.89 |
| RW_Kirehe_Bovine_2025_29 | Bovine | 2025 | Kirehe | N/A | WGS | 66.31 | 92.07 | 96.06 |

**Table footnotes:** S; Small, M; Medium, L; Large, Seg; segment, WGS; whole-genome sequencing. \*coverage depth set at x6.

We were able to generate 56 RVFV segment genomes (≥80% genome coverage and 6x depth) for subsequent phylogenetic analysis using a rapidly designed amplicon assay. Using the Nextstrain pipeline, we generated three timetree phylogenies for each RVFV segment (Figures 2a-f). All our obtained sequences formed a cluster within lineage C, uniquely with sequence data from Rwanda, with a most recent common ancestor (MRCA) estimated to originate from 2021 (95% confidence intervals) (S and L segments) and 2022 (95% confidence intervals) (M segment). This suggests that 2024 and 2025 cases are a consequence of a single introduction in the region and continued local spread. Indeed, the 2024 outbreak primarily occurred within Rwanda’s Eastern Province, a region highly predisposed for transmission, with several lakes and low altitudes ideal for mosquito proliferation (12). Meanwhile, sequences from 2025 formed more sporadic regional clusters within the 2022 outbreak, distinct from the 2024 outbreak, additionally suggesting low-level local persistence. This pattern is consistent with sustained circulation of the original outbreak lineage, which is estimated to have emerged in mid-2021 based on the MRCA analysis. Whether maintenance of virus circulation involves undetected presence in livestock or in wild animal reservoirs remains to be determined. Periodic spillover events may have resulted in distinct epidemic clusters or transmission chains, which could explain why the 2024 and 2025 sequences cluster separately and branch from the 2022 lineage, rather than forming a shared monophyletic group. A caveat is that the interpretation of phylogenetic analysis is critically dependent on the availability of sequence data from the wider region, and for instance we cannot rule out new cross border introductions. RVFV sequences isolated within Rwanda between 2022-2025 also showed a close phylogenetic relationship with those isolated from Uganda in 2023 and additional sequences from neighbouring countries likely would reveal further relationships. Moreover our obtained sequences did not cluster with any vaccine strains within lineage E, comparatively to the 2022 outbreak, or with the recently declared 2025 outbreaks in Senegal and Mauritania within lineage H (13).

**Figure 2.**
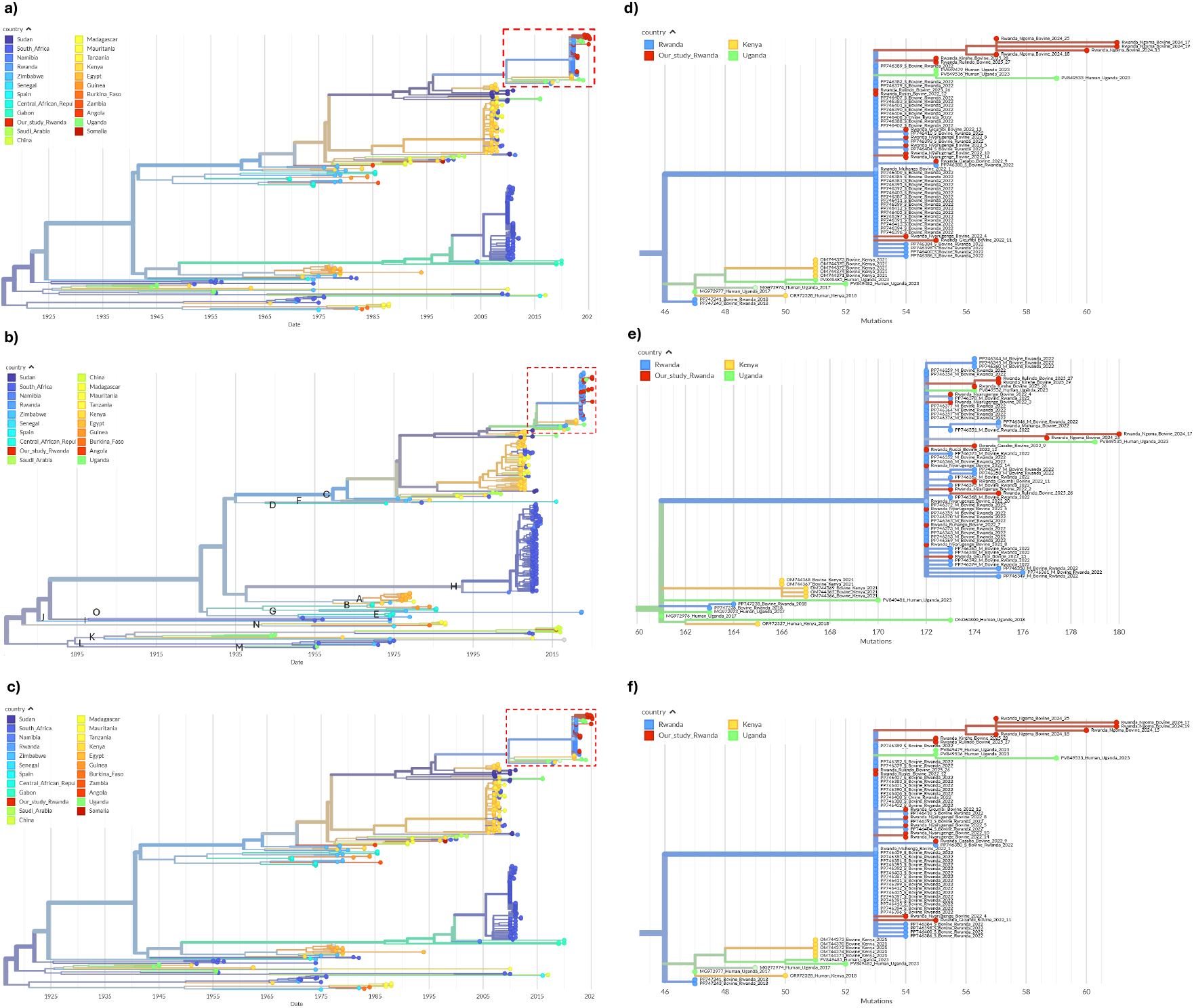
Phylogenetic analysis was performed using Nextstrain for all RVFV segments. Reference sequences were downloaded from the BV-BRC database (https://www.bv-brc.org/), along with sequences from the 2022 outbreak in Rwanda (7). The rectangular dotted red box represents the cluster including our consensus sequences for each segment: (a) Segment S, (b) Segment M, (c) Segment L. We have also provided an enlarged view of our highlighted cluster providing higher resolution of divergence: (d) Segment S enlarged (e) Segment M enlarged (f) Segment L enlarged.

Low genomic diversity has been observed for RVFV with studies indicating the virus has a limited tolerance for mutations (14). Overall, the observed substitution rate was 2.65×10^−4^ (0.44 subs/genome/year), 2.55×10^−4^ (0.98 subs/genome/year) and 2.47×10^−4^ (1.59 subs/genome/year) for S, M and L segments respectively. These rates are similar to previously reported rates of 5.1×10^−4^ (S segment), 2.7×10^−4^ (M segment), 1.7×10^−4^ (L segment) from Nsengimana and colleagues (7). Sustained genomic surveillance and variation detection in endemic areas is essential to monitor vaccine efficiency and ensure the specificity of diagnostic real-time PCR assays. This is particularly important as vaccine campaigns and testing tend to be more reactive in response to growing outbreaks, rather than proactive nationwide preventative strategies (15).

Future studies should not only assess vaccine coverage, but also evaluate post-vaccination immunity and monitor breakthrough infections.

This study had several limitations. Only a limited number of samples were available, and more representative sampling, systematic collection, sample storage, as well as vaccine status of the animals could improve our interpretation. As assay development was initiated in response to rising RVFV cases ahead of the workshop, time constraints precluded comprehensive amplicon validation and primer balancing, resulting in low coverage in two genome segments. Nevertheless, our study provides a proof-of-principle approach which can be applied to other known and unknown infectious viruses within low-and-middle-income countries.

## Conclusion

RVFVs circulating in 2024 and 2025 in Rwanda were closely related to viruses detected in 2022, providing evidence for ongoing circulation rather than the emergence of a novel strain. Region specific clustering could provide evidence of ongoing silent circulation within naïve livestock populations or wildlife reservoirs, underpinning the critical importance of genomic surveillance at the wildlife–livestock interface. More genomic surveillance from East and Sub-Saharan Africa is crucial to increase outbreak resolution. Strengthening surveillance, investing in research and addressing environmental triggers are crucial for future preparedness.

## Supporting information

Supplementary Materials_Metagenomics sequencing

## Funding and Acknowledgements

This work was funded by the Global Health EDCTP3 Joint Undertaking (Global Health EDCTP3) programme under grant agreement No. 101103059 (GREATLIFE) and 101195116 (JUA KIVU). We greatly thank the Rwanda Agriculture and Animal Resources Development Board, Kigali, Rwanda, for their collaboration during the study. Finally, we also acknowledge the EuFMD 10th FAR Funding Program for the research support they awarded to JCU, FN and PN.

## Notes

### Competing Interest Statement

The authors have declared no competing interest.

