## Supplementary Materials_Metagenomics sequencing for "Persistent circulation of Rift Valley fever virus lineage C in Rwanda, 2022–2025"

##### *Metagenomics sequencing*

Sequencing libraries were prepared using the Native Barcoding Kit 24 v14 (SQK-NBD114.24) (Oxford Nanopore Technologies) and sequenced on a R.10.4.1 MinION flowcell. Sequencing reads were trimmed by fastp v0.20.1 (https://github.com/OpenGene/fastp) and cutadapt v3.0 (https://github.com/marcelm/cutadapt). Minimap2 v2.28 (https://github.com/lh3/minimap2) was used to remove host reads, followed by taxonomic classification with KRAKEN2 v.2.17.1 (https://github.com/jenniferlu717/KrakenTools) and further confirmation using NCBI BLAST. Viral consensus sequences were generated with Minimap2 and Samtools v1.12 (https://github.com/samtools).

##### *Amplicon sequencing*

Random priming was first performed using 50µM random hexamers, 10mM dNTP and 6.5 µl of extracted RNA and incubated at 65°C for 5 min followed by 4°C for 2 min. cDNA was then synthesized using SuperScript IV reverse transcriptase, superscript IV buffer (5x) and DTT (100mM) (Invitrogen, Carlsbad, CA, USA) and incubated at 23°C for 10 min, 50°C for 60 min and then 80°C for 10 min. Amplicons were then generated using the Q5® High-Fidelity DNA Polymerase kit, dNTP mix (10nM) and each respective primer pool (10µM) (Integrated DNA Technologies, Coralville, IA, USA) (Table S2). PCR conditions were 98°C for 45 s, followed by 35 cycles of 98°C for 15 s and 65°C for 5 min. Sequencing libraries were generated using 100-150ng as input for the Native Barcoding Kit 24 v14 (SQK-NBD114.24) (ONT) and sequenced on a R10.4.1 flowcell (FLO-MIN114) (ONT). Primer sequences were trimmed by Ivar v1.4.4 (https://github.com/andersen-lab/ivar) followed by read trimming using fastp v0.20.1.

Consensus sequences were generated using MiniMap2 and an inhouse reference database (Table S1). Phylogenetic analysis was performed using augur v28.0 (https://github.com/augurproject) on Nextstrain (https://github.com/augurproject). The RVFV typing tool was used for typing the glycoprotein gene (Gn) (M segment). RVFV genome sequences were deposited onto GenBank.

### Supplementary Tables and Figures

#### *Table S1. GenBank accession numbers of all reference sequences used in Nextstrain analysis*

| S segment references | M segment references | L segment references |
|---|---|---|
| PP746382 | PP746342 | PP746304 |
| PP746399 | PP746354 | PP746336 |
| PP746412 | PP746369 | PP746315 |
| PP746387 | PP746364 | PP746322 |
| PP746400 | PP746376 | PP746337 |
| PP746403 | PP746374 | PP746326 |
| PP746383 | PP746365 | PP746328 |
| PP746389 | PP746353 | PP746329 |
| PP746379 | PP746352 | PP746313 |
| PP746395 | PP746366 | PP746324 |
| PP746401 | PP746355 | PP746335 |
| PP746402 | PP746378 | PP746317 |
| PP746407 | PP746368 | PP746311 |
| PP746410 | PP746351 | PP746334 |
| PP746386 | PP746371 | PP746318 |
| PP746384 | PP746373 | PP746325 |
| PP746398 | PP746345 | PP746314 |
| PP746404 | PP746347 | PP746340 |
| PP746381 | PP746372 | PP746316 |
| PP746409 | PP746348 | PP746339 |
| PP746411 | PP746358 | PP746309 |
| PP746392 | PP746360 | PP746305 |
| PP746393 | PP746343 | PP746307 |
| PP746380 | PP746362 | PP746308 |
| PP746408 | PP746344 | PP746306 |
| JQ820482 | PP746375 | PP746327 |
| EU574057 | PP746349 | PP746310 |
| EU574058 | PP746361 | PP746338 |
| EU574059 | PP746346 | PP746330 |
| EU574060 | PP746350 | PP746323 |
| EU574061 | EU574042 | MT561464 |
| EU574062 | EU574041 | MT561461 |
| EU574063 | EU574044 | OM744375 |
| EU574064 | EU574043 | OM744376 |
| EU574065 | EU574045 | OM744377 |
| EU574066 | EU574046 | OM744378 |
| EU574067 | EU574047 | OM744379 |
| EU574068 | EU574048 | JQ820483 |
| EU574069 | EU574049 | MG659820 |
| EU574070 | EU574050 | MG659819 |
| EU574071 | EU574051 | MG659821 |
| EU574072 | EU574052 | MG659822 |
| EU574073 | EU574053 | MG659823 |
| EU574074 | EU574054 | MG659824 |
| EU574075 | EU574055 | MG659825 |
| EU574076 | EU574056 | MG659826 |
| EU574077 | MT561460 | MG659827 |
| EU574078 | OM744365 | MG659828 |
| EU574079 | OM744366 | MG659829 |
| EU574080 | OM744367 | MG659831 |
| EU574081 | OM744369 | MG659830 |
| EU574082 | OM744368 | MG659832 |
| EU574083 | MG659733 | MG659834 |
| EU574084 | MG659734 | MG659833 |
| EU574085 | MG659735 | MG659835 |
| EU574086 | MG659736 | MG659836 |
| EU574087 | MG659737 | MG659837 |
| EU709747 | MG659738 | MG659838 |
| EU709748 | MG659739 | MG659839 |
| JQ820472 | MG659740 | MG659840 |
| JQ820474 | MG659741 | MG659841 |
| JQ820473 | MG659742 | MG659842 |
| JQ820475 | MG659743 | MG659843 |
| JQ820476 | MG659744 | MG659844 |
| MT561462 | MG659745 | MG659845 |

| | | |
|---|---|---|
| MT561459 | MG659747 | MG659846 |
| OM744370 | MG659746 | MG659847 |
| OM744371 | MG659748 | MG659849 |
| OM744372 | MG659749 | MG659848 |
| OM744373 | MG659750 | JQ820484 |
| OM744374 | MG659752 | MG659850 |
| KU167025 | MG659751 | MG659852 |
| MG659907 | MG659753 | MG659853 |
| MG659908 | MG659754 | MG659851 |
| MG659905 | MG659756 | MG659854 |
| MG659906 | MG659755 | MG659855 |
| MG659909 | MG659757 | MG659856 |
| MG659910 | MG659758 | MG659857 |
| MG659911 | MG659759 | MG659858 |
| MG659912 | MG659760 | MG659859 |
| MG659913 | MG659761 | MG659860 |
| MG659914 | MG659762 | MG659861 |
| MG659916 | MG659763 | MG659862 |
| MG659915 | MG659764 | MG659863 |
| MG659917 | MG659765 | MG659864 |
| MG659918 | MG659766 | MG659865 |
| MG659919 | MG659767 | MG659866 |
| MG659920 | MG659768 | MG659868 |
| MG659921 | MG659770 | MG659867 |
| MG659922 | MG659771 | MG659869 |
| MG659923 | MG659769 | MG659870 |
| MG659924 | MG659772 | MG659871 |
| MG659925 | MG659773 | MG659872 |
| MG659926 | MG659774 | MG659873 |
| MG659927 | MG659775 | MG659874 |
| MG659928 | MG659777 | MG659875 |
| MG659929 | MG659776 | MG659876 |
| MG659930 | MG659778 | MG659877 |
| MG659931 | MG659779 | MG659878 |
| MG659932 | MG659780 | MG659879 |
| MG659933 | MG659782 | MG659880 |
| MG659934 | MG659781 | MG659882 |
| MG659935 | MG659783 | MG659881 |
| MG659936 | MG659784 | MG659883 |
| MG659937 | MG659785 | MG659884 |
| MG659938 | MG659786 | MG659885 |
| MG659939 | MG659787 | MG659886 |
| MG659940 | MG659788 | MG659887 |
| MG659941 | MG659789 | MG659888 |
| MG659942 | MG659790 | MG659889 |
| MG659943 | MG659792 | MG659890 |
| MG659944 | MG659791 | MG659891 |
| MG659946 | MG659793 | MG659892 |
| MG659945 | MG659794 | MG659893 |
| MG659947 | MG659795 | MG659894 |
| MG659948 | MG659796 | MG659895 |
| MG659949 | MG659798 | MG659896 |
| MG659950 | MG659799 | MG659897 |
| MG659951 | MG659800 | MG659898 |
| MG659952 | MG659801 | MG659899 |
| MG659953 | MG659802 | MG659900 |
| MG659954 | MG659803 | MG659901 |
| MG659955 | MG659797 | MG659902 |
| MG659956 | MG659804 | MG659904 |
| MG659957 | MG659805 | MG659903 |
| MG659958 | MG659806 | KU167027 |
| MG659959 | MG659807 | JQ820485 |
| MG659960 | MG659810 | MZ513011 |
| MG659961 | MG659808 | MZ513012 |
| MG659962 | MG659809 | MZ513013 |
| MG659963 | MG659811 | KY126665 |
| MG659965 | MG659812 | KY126666 |
| MG659964 | MG659813 | KY126668 |
| MG659966 | MG659814 | KY126667 |
| MG659967 | MG659815 | KY126669 |
| MG659968 | MG659816 | KY126671 |

| | | |
|---|---|---|
| MG659969 | MG659817 | KY126670 |
| MG659971 | MG659818 | KY126672 |
| MG659970 | KU167026 | KY126673 |
| MG659972 | KY126690 | KY126674 |
| MG659973 | KY126691 | KY126675 |
| MG659974 | KY126692 | KY126677 |
| MG659975 | KY126693 | KY126676 |
| MG659976 | KY126694 | KY126678 |
| MG659977 | KY126696 | KY126679 |
| MG659978 | KY126695 | KY126680 |
| MG659979 | KY126698 | KY126681 |
| MG659980 | KY126697 | KY126683 |
| MG659981 | KY126700 | KY126682 |
| MG659982 | KY126699 | KY126684 |
| MG659983 | KY126701 | KY126685 |
| MG659984 | KY126702 | KY126686 |
| MG659986 | KY126703 | KY126687 |
| MG659985 | KY126704 | KY126688 |
| MG659987 | KY126705 | KY126689 |
| MG659988 | KY126706 | JQ820486 |
| MG659990 | KY126707 | OM744403 |
| MG659989 | KY126708 | OP146107 |
| KY196487 | KY126709 | OP146110 |
| KY196488 | KY126710 | OR972326 |
| KY196489 | KY126711 | MG953422 |
| KY196490 | KY126712 | MG953423 |
| KY196492 | KY126713 | MG972972 |
| KY196491 | KY126714 | MG972975 |
| KY196493 | OM744404 | MG972978 |
| KY196494 | ON060800 | KU978779 |
| KY196495 | JQ820487 | KU978780 |
| KY196496 | OP146105 | KU978781 |
| KY196497 | OP146108 | KX096938 |
| KY196498 | OP146111 | KX096941 |
| KY196499 | OR125122 | KX609031 |
| KY196500 | MG953419 | KX611605 |
| KY196501 | OR972327 | KX632066 |
| KY196502 | JQ820488 | KJ782454 |
| KY196503 | JQ820489 | KJ782457 |
| KY196504 | MG972973 | KX944849 |
| KY196505 | JQ820490 | KX944850 |
| KY196506 | MG972976 | KX944851 |
| KY196508 | JQ820491 | KX944852 |
| KY196509 | KU978776 | KX944853 |
| KY196510 | KU978778 | KX944854 |
| KY196511 | KX096939 | KX944855 |
| KY196507 | KX096942 | KX944856 |
| OM744405 | KX609032 | KX944857 |
| OP146106 | KX611606 | KX944858 |
| OP146109 | KX632067 | KX944859 |
| OP146112 | KJ782453 | KX944860 |
| OR972328 | KJ782456 | KX944861 |
| OR528950 | KX944826 | KX944862 |
| OR528951 | KX944827 | KX944863 |
| OR528953 | KX944828 | KX944864 |
| OR528952 | KX944829 | KX944865 |
| MG953425 | KX944830 | KX944866 |
| MG953426 | KX944831 | KX944868 |
| OR805807 | KX944833 | KX944869 |
| MG972974 | KM210509 | MG649050 |
| MG972977 | KX944834 | KX944870 |
| JQ840745 | KX944835 | KX944871 |
| JQ840746 | KX944836 | HM586953 |
| KU978773 | KX944837 | HM586954 |
| KX096940 | KX944838 | HM586955 |
| KX096943 | KX944839 | HM586956 |
| KX611607 | KX944840 | HM586957 |
| KJ782452 | KX944841 | HM586958 |
| KJ782455 | KX944842 | HM586959 |
| KX944803 | KX944843 | HM586960 |
| KX944804 | KX944844 | MG649051 |

| | | |
|---|---|---|
| KX944805 | KX944845 | HM586961 |
| KX944806 | KX944846 | HM586962 |
| KX944807 | KX944847 | HM586963 |
| KX944808 | KX944848 | MG649052 |
| KX944809 | HM586964 | MG649053 |
| KX944810 | HM586965 | DQ375395 |
| KX944811 | HM586966 | DQ375396 |
| KX944812 | HM586967 | DQ375397 |
| KX944813 | HM586968 | DQ375398 |
| KX944816 | HM586969 | DQ375399 |
| KX944817 | HM586970 | DQ375400 |
| KX944818 | HM586971 | DQ375401 |
| KX944819 | HM586972 | DQ375402 |
| KX944820 | HM586973 | DQ375403 |
| KX944821 | HM586974 | DQ375404 |
| KX944822 | MG649054 | DQ375405 |
| KX944823 | MG649055 | DQ375406 |
| KX944825 | MG649056 | DQ375407 |
| KM210508 | MG649057 | DQ375408 |
| JQ820477 | NC_014396 | DQ375409 |
| HM586975 | HQ009512 | DQ375410 |
| HM586976 | DQ380183 | DQ375411 |
| HM586977 | DQ380184 | DQ375412 |
| HM586978 | DQ380185 | DQ375413 |
| HM586979 | DQ380186 | DQ375414 |
| HM586980 | DQ380187 | DQ375415 |
| HM586981 | DQ380188 | DQ375416 |
| HM586982 | DQ380189 | DQ375417 |
| HM586983 | DQ380191 | DQ375418 |
| HM586984 | DQ380194 | NC_014397 |
| JQ820478 | DQ380195 | DQ375419 |
| NC_014395 | DQ380196 | DQ375420 |
| DQ380143 | DQ380197 | DQ375421 |
| DQ380144 | DQ380198 | DQ375422 |
| DQ380145 | DQ380199 | DQ375423 |
| DQ380146 | DQ380200 | DQ375424 |
| DQ380149 | DQ380201 | DQ375426 |
| DQ380151 | DQ380202 | DQ375425 |
| DQ380152 | DQ380203 | DQ375428 |
| DQ380153 | DQ380204 | DQ375429 |
| DQ380154 | DQ380205 | DQ375430 |
| DQ380155 | DQ380206 | DQ375431 |
| DQ380156 | DQ380207 | DQ375432 |
| DQ380157 | DQ380209 | DQ375433 |
| DQ380158 | DQ380210 | DQ375434 |
| DQ380159 | DQ380211 | JF311368 |
| DQ380160 | DQ380212 | JF311369 |
| DQ380161 | DQ380214 | JF311370 |
| DQ380162 | DQ380215 | JF311371 |
| DQ380163 | DQ380216 | JF311372 |
| DQ380164 | DQ380217 | JF311373 |
| DQ380165 | DQ380218 | JF311374 |
| DQ380166 | DQ380219 | JF311375 |
| DQ380167 | DQ380220 | JF311376 |
| DQ380168 | DQ380221 | JF326186 |
| DQ380169 | DQ380222 | JF326187 |
| DQ380170 | EF460404 | JF326188 |
| DQ380171 | EF467177 | JF326189 |
| DQ380172 | EF467178 | JF326190 |
| DQ380173 | JF309200 | JF784386 |
| DQ380174 | JF311377 | JQ068144 |
| DQ380175 | JF311378 | EU574004 |
| DQ380177 | JF311379 | EU574005 |
| DQ380178 | JF311380 | EU574006 |
| DQ380179 | JF311381 | EU574007 |
| DQ380180 | JF311382 | EU574008 |
| DQ380181 | JF311383 | EU574009 |
| JQ820479 | JF311384 | EU574010 |
| JF311386 | JF311385 | EU574011 |
| JF311387 | JF326191 | EU574012 |
| JF311388 | JF326192 | EU574013 |

| | | |
|---|---|---|
| JF311389 | JF326193 | EU574014 |
| JF311390 | JF326194 | EU574015 |
| JF311391 | JF326195 | EU574016 |
| JF311392 | JF784387 | EU574017 |
| JF311393 | JQ068143 | EU574018 |
| JF311394 | EU574031 | EU574019 |
| JQ820480 | EU574032 | EU574020 |
| JF326196 | EU574033 | EU574021 |
| JF326197 | EU574034 | EU574022 |
| JF326198 | EU574035 | EU574023 |
| JF326199 | EU574036 | EU574024 |
| JF326200 | EU574037 | EU574025 |
| JF326201 | EU574038 | EU574026 |
| JF326202 | EU574039 | EU574027 |
| JF326203 | PP747236 | EU574028 |
| JF326204 | PP747238 | EU574029 |
| JF784388 | DQ380213 | EU574030 |
| EU312103 | DQ380193 | PP747231 |
| EU312104 | PP746359 | PP747233 |
| EU312105 | PP746363 | OP146104 |
| EU312106 | PP746357 | PP746321 |
| EU312107 | PP746367 | PP746331 |
| EU312108 | PP746370 | PP746332 |
| EU312109 | PP746377 | PP746333 |
| EU312110 | PP746356 | PP746312 |
| EU312111 | PV849481 | PP746319 |
| EU312112 | PV849532 | PP746320 |
| EU312113 | PV849535 | PV849534 |
| EU312114 | | PV849531 |
| EU312115 | | PV849480 |
| EU312116 | | |
| EU312117 | | |
| EU312118 | | |
| EU312119 | | |
| EU312120 | | |
| EU312121 | | |
| EU312122 | | |
| EU312123 | | |
| EU312124 | | |
| EU312125 | | |
| EU312126 | | |
| EU312127 | | |
| EU312128 | | |
| EU312130 | | |
| EU312131 | | |
| EU312132 | | |
| EU312133 | | |
| EU312134 | | |
| EU312135 | | |
| EU312136 | | |
| EU312137 | | |
| EU312138 | | |
| EU312139 | | |
| EU312140 | | |
| EU312141 | | |
| EU312142 | | |
| EU312143 | | |
| EU312144 | | |
| EU312145 | | |
| EU312146 | | |
| EU312147 | | |
| JQ820481 | | |
| PP747241 | | |
| PP747243 | | |
| DQ380182 | | |
| PP746388 | | |
| PP746397 | | |
| PP746406 | | |
| PP746390 | | |
| PP746394 | | |
| PP746396 | | |

| |
|---|
| PP746405 |
| PP746413 |
| PP746385 |
| PP746391 |
| PV849536 |
| PV849533 |
| PV849485 |
| PV849482 |
| PV849479 |

*Table S2. RVFV Amplicon primers*

| Name | Position | Position | Pool | Sense | Sequence |
|---|---|---|---|---|---|
| SegL_RVFV_1_A | 22 | 43 | 1 | + | ACACAAAGGCGCCCAATCATG |
| SegL_RVFV_2_A | 567 | 590 | 1 | - | TGCTTCAGAGTCCTCTAGCTCCA |
| SegL_RVFV_3_B | 459 | 487 | 2 | + | CCATGACTAAACTCGCTAAGTATGAGGT |
| SegL_RVFV_4_B | 1009 | 1031 | 2 | - | CCAGGGAGGCAGCTGAATAGTT |
| SegL_RVFV_5_A | 890 | 920 | 1 | + | GGGAATGACAAAGTTCTGAGATTTTCAAAA |
| SegL_RVFV_6_A | 1440 | 1471 | 1 | - | TCTTTATCACAAGAGAGAGGGTTGTATAGAT |
| SegL_RVFV_7_B | 1333 | 1360 | 2 | + | GCAAATGAGAAATAGGAGTAAGCAGCC |
| SegL_RVFV_8_B | 1883 | 1906 | 2 | - | TCTGCCCAGAATGAGAAGGAGGA |
| SegL_RVFV_9_A | 1777 | 1802 | 1 | + | GGTCTTCAAGCCCTACATAGATGCT |
| SegL_RVFV_10_A | 2327 | 2350 | 1 | - | GGCTCGGTCTCCTCTTCTTTGTT |
| SegL_RVFV_11_B | 2210 | 2243 | 2 | + | ATTCTTAAGAAAAAGGATGGATCTATATCCTGG |
| SegL_RVFV_12_B | 2760 | 2786 | 2 | - | ACAAATATGCATGCATCCTTGTGACT |
| SegL_RVFV_13_A | 2651 | 2678 | 1 | + | GCAGACAAGAACTACACAAGGGATAAA |
| SegL_RVFV_14_A | 3201 | 3231 | 1 | - | CATGATAAGCTTTGAAGAGATCCATCACAA |
| SegL_RVFV_15_B | 3094 | 3119 | 2 | + | GCTGATCATTAGGGGATGCTCAATG |
| SegL_RVFV_16_B | 3644 | 3667 | 2 | - | GCTGCAATCCATCTGATCGTTGG |
| SegL_RVFV_17_A | 3540 | 3564 | 1 | + | TGGGAGTGTACCTTGCCATTTACC |
| SegL_RVFV_18_A | 4090 | 4118 | 1 | - | CCTAGATCCCCACTTTAGAGAAGAGCTA |
| SegL_RVFV_19_B | 3925 | 3957 | 2 | + | AGGTTTCAGATTTAATCTCTTCAAAGCTATCA |
| SegL_RVFV_20_B | 4475 | 4505 | 2 | - | ATCCAGTGATTCTAATTCCTCAATATTCGG |
| SegL_RVFV_21_A | 4362 | 4391 | 1 | + | CTGAGTTCAACTTCTTGGAGGATTCTAAG |
| SegL_RVFV_22_A | 4912 | 4934 | 1 | - | AGCAAGGTCTTCTACACCTCGG |
| SegL_RVFV_23_B | 4806 | 4831 | 2 | + | GCAAGCCCAGAACAGTCAGAATAAC |
| SegL_RVFV_24_B | 5356 | 5380 | 2 | - | GCCCACTGCCTTATTGGAACACTC |
| SegL_RVFV_25_A | 5249 | 5272 | 1 | + | GGCTTCATGGACGGATATCAGGT |
| SegL_RVFV_26_A | 5799 | 5830 | 1 | - | AGTCTGGTTCTGTTAATTCCTTTTATCTGTC |
| SegL_RVFV_27_B | 5692 | 5717 | 2 | + | TCAAGCCCTGCAATTATTTGAGAGG |
| SegL_RVFV_28_B | 6242 | 6267 | 2 | - | TCCTGATGCTTTCTTCTGGTCTCTG |
| SegL_RVFV_29_A | 5847 | 5876 | 1 | + | TTTGTTCTGAATCTTCCCTGAGATCAAAA |
| SegL_RVFV_30_A | 6397 | 6423 | 1 | - | GACCGTCCAATATTGTAGCACTATGC |
| SegM_RVFV_1_A | 0 | 28 | 1 | + | ACACAAAGACGGTGCATTAAATGTATGT |

| | | | | | |
|---|---|---|---|---|---|
| SegM_RVFV_2_A | 525 | 547 | 1 | - | TCCTCCTGAGTCATCCCGTCAA |
| SegM_RVFV_3_B | 383 | 410 | 2 | + | GATGCTAAGCAAATAGGGAGAGAAACC |
| SegM_RVFV_4_B | 932 | 959 | 2 | - | TTTTGATCCATCATCCTCACTTGACTG |
| SegM_RVFV_5_A | 812 | 835 | 1 | + | AAAATGGCTTCAGTCAAGTGCCC |
| SegM_RVFV_6_A | 1362 | 1384 | 1 | - | AAAGGCTTCTTCCAGACCCCTG |
| SegM_RVFV_7_A | 1362 | 1388 | 1 | - | ACATAAAGGTTTCTTCCAGACCCCTG |
| SegM_RVFV_8_B | 1198 | 1226 | 2 | + | GCTCAAAAGCTTTGATATCTCTCAGTGC |
| SegM_RVFV_9_B | 1736 | 1762 | 2 | - | GTGTGACACTGGTAATTTATCAGGCC |
| SegM_RVFV_10_A | 1568 | 1599 | 1 | + | AGTCCTTCTACTGAGATTACACTCAAGTATC |
| SegM_RVFV_11_A | 1570 | 1599 | 1 | + | TCCTTCTACCGAGATTACACTCAAGTATC |
| SegM_RVFV_12_A | 2113 | 2136 | 1 | - | TGGAGCAAGTGGTGATTCTGGAG |
| SegM_RVFV_13_B | 1973 | 1995 | 2 | + | TGGATGGAAGGAGGTCAGTTGG |
| SegM_RVFV_14_B | 2513 | 2538 | 2 | - | GCAGATACGTGTGCACAAATAAGCA |
| SegM_RVFV_15_A | 2406 | 2433 | 1 | + | CTTCAGCAGAGTTTTCATTTGTTGGAG |
| SegM_RVFV_16_A | 2408 | 2433 | 1 | + | TCAGCAGAGTTTTCATTTGTTGGGG |
| SegM_RVFV_17_A | 2956 | 2980 | 1 | - | TCATTCCTTGTCTGTGGCAGAGAA |
| SegM_RVFV_18_B | 2839 | 2862 | 2 | + | CCTGAGTGCTCATGAATCATGCC |
| SegM_RVFV_19_B | 3358 | 3388 | 2 | - | CTATCATCAAATGGGTCAATAGCTATCAGG |
| SegM_RVFV_20_B | 3358 | 3390 | 2 | - | GCCTATCATCAAATGGATCAATAGCTATCAGA |
| SegM_RVFV_21_A | 3207 | 3230 | 1 | + | ACAATAAGGACGGGTCTCTGCAT |
| SegM_RVFV_22_A | 3205 | 3230 | 1 | + | CCACAATAAGGATGGGTCTCTGCAT |
| SegM_RVFV_23_A | 3751 | 3774 | 1 | - | CCCCACCACCCCAAATAACAACT |
| SegM_RVFV_24_A | 3751 | 3774 | 1 | - | CCCCACCACCCCAAATTACAACT |
| SegM_RVFV_25_B | 3400 | 3424 | 2 | + | GGGGGAATCAACAGTTGTGAATCC |
| SegM_RVFV_26_B | 3398 | 3424 | 2 | + | GGGGGAGAATCAATAGTTGTGAATCC |
| SegM_RVFV_27_B | 3863 | 3885 | 2 | - | ACACAAAGACCGGTGCAACTTC |
| SegS_RVFV_1_A | 0 | 24 | 1 | + | ACACAAAGACCCCCTAGTGCTTAT |
| SegS_RVFV_2_A | 524 | 548 | 1 | - | GGATAGCCTCGGTCACTATCATCC |
| SegS_RVFV_3_B | 378 | 400 | 2 | + | TGGAGAACCCTCACTGGCTTTC |
| SegS_RVFV_4_B | 928 | 950 | 2 | - | GCTGCTCAGGCCTACAAAACAG |
| SegS_RVFV_5_A | 825 | 848 | 1 | + | TGATTAGAGGTTAAGGCTGCCCC |
| SegS_RVFV_6_A | 1375 | 1403 | 1 | - | CTCATCAACAAGTACAAGCTAAAGGAGG |
| SegS_RVFV_7_B | 1125 | 1147 | 2 | + | CGGGAGAACTGCAGCAGATACA |
| SegS_RVFV_8_B | 1635 | 1664 | 2 | - | ACAATAATGGACAACTATCAAGAGCTTGC |

***Table footnotes:*** *SegL; Segment L, SegM; Segment M, SegS; Segment S, RVFV; Rift Valley fever virus*

*Table S3. Overview of RVFV cases in Rwanda between 2022-2025*

| Districts tested | Collection period | Positive districts | Total no. of animal samples tested | Total no. of positives | Species | Case fatality | Percent positivity | Refs |
|---|---|---|---|---|---|---|---|---|
| All (30 districts) | March - December 2022 | Muhanga, Nyarugenge ,Ruhango, Gasabo, Gicumbi, Rusizi | 1,880,591 | 1342 | Cattle (95.8%), goats (2.5%) and sheep (1.7%) | 38% | 0.07% | 1, 2 |
| Gicumbi | October 2023 | Gicumbi | 1288 | 5 | N/A | N/A | 0.38% | * |
| Gasabo, Kamonyi, Nyarugenge, Musanze, Gatsibo, Kirehe and Ngoma* | March – October 2024 | Ngoma | 2692 | 22 | Cattle (45.8%), goats (33.3%) and sheep (20.8%) | 8.30% | 0.90% | 3 |
| Muhanga, Rulindo and Kirehe | March - April and October 2025 | Rulindo and Kirehe | 936 | 18 | N/A | N/A | 1.92% | * |

***Table footnotes:*** *N/A; not available. * Personal communication from the Chief Veterinary Officer, National Veterinary Reference Laboratory in Rubirizi, Rwanda, December 2025*

*References:*

1. Remera E, Rwagasore E, Nsekuye O, Semakula M, Gashegu M, Rutayisire R, Ishema L, Musanabaganwa C, Butera Y, Nsanzimana S, Muvunyi CM, Ahmed A. Rift Valley Fever Epizootic, Rwanda, 2022. Emerg Infect Dis. 2024 Oct;30(10):2191-2193. doi: 10.3201/eid3010.240264. PMID: 39320366; PMCID: PMC11431898.
2. Ishema L, Colombe S, Ndayisenga F, Uwibambe E, Van Damme E, Meudec M, Rwagasore E, Mugwaneza D, Van Bortel W, Shyaka A. One Health investigation and response to a nationwide outbreak of Rift Valley fever in Rwanda - March to December 2022. One Health. 2024 Jul 4;19:100854. doi: 10.1016/j.onehlt.2024.100854. PMID: 40206839; PMCID: PMC11980628.
3. Ministry of Agriculture and Animal Resources (Rwanda). Rwanda tackles RVF using 'One Health' approach. MINAGRI. Published 26 September 2024. Available at: https://www.minagri.gov.rw/updates/news-details/rwanda-tackles-rvf-using-one-health-approach (accessed 20/01/2026)

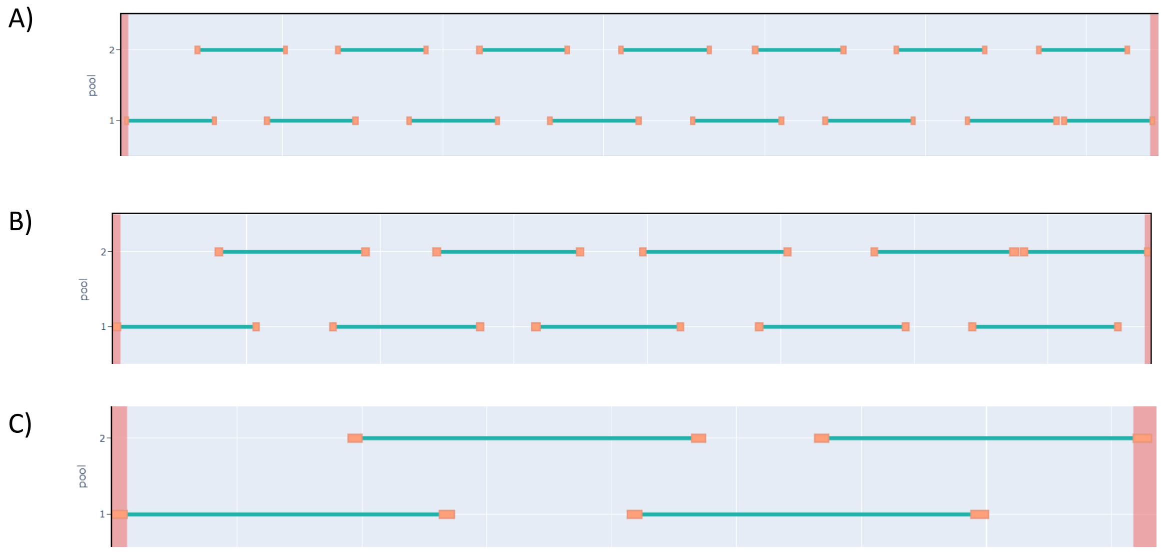

*Figure S1. Rift Valley fever virus Primer overlapping regions. A) Segment L, B) Segment M, C) Segment S*

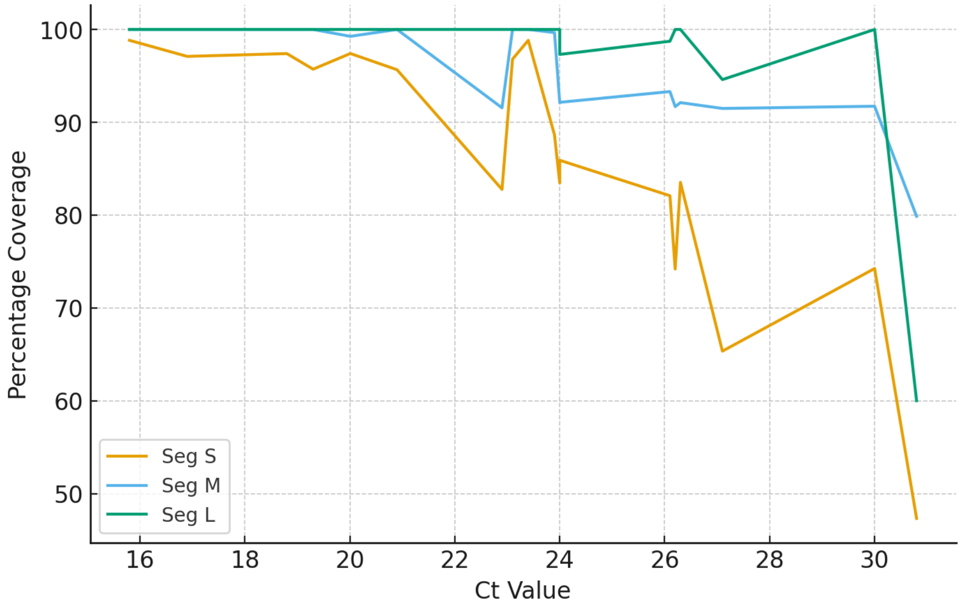

*Figure S2. Ct values compared to percentage coverage*

A)

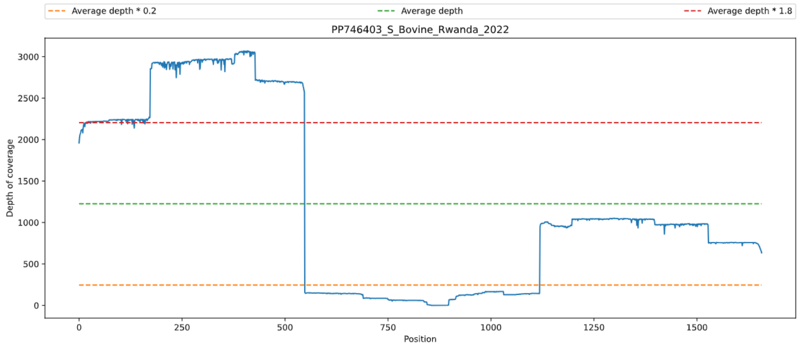

B)

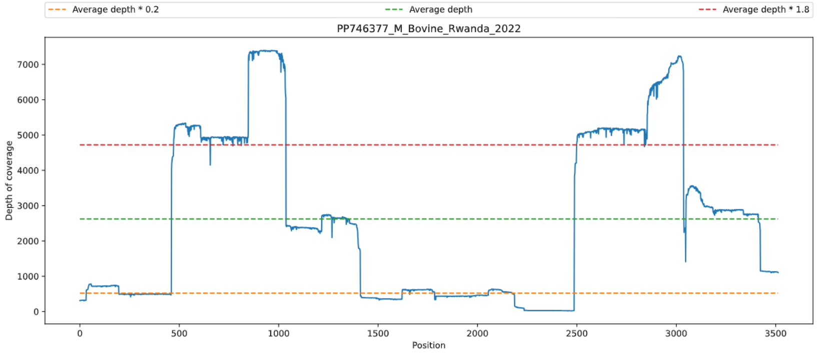

C)

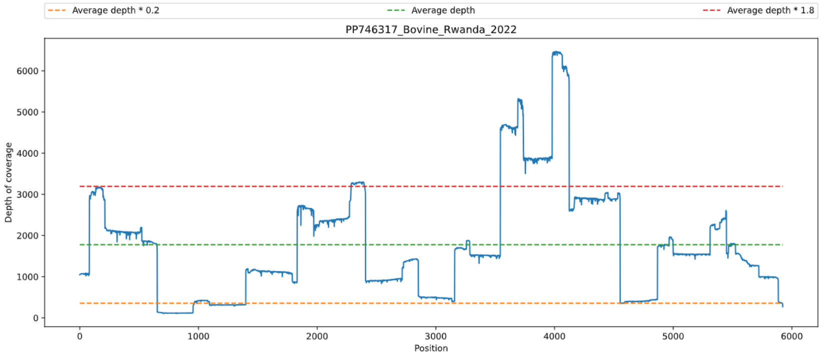

*Figure S3. Example coverage patterns for amplicon sequencing. A) Segment S, B) Segment M, C) Segment L. Two amplicon dropouts can be observed within Segment M (299bps at approx. position 2188-2487) and in Segment S (570 bps at approx. position 549-1119).*
